# The functional and structural characterization of *Xanthomonas campestris* pv. *campestris* core effector XopP revealed a new kinase activity

**DOI:** 10.1101/2022.10.27.514000

**Authors:** Konstantinos Kotsaridis, Vassiliki A. Michalopoulou, Dimitra Tsakiri, Dina Kotsifaki, Aikaterini Kefala, Nikolaos Koundourakis, Patrick H.N. Celie, Michael Kokkinidis, Panagiotis F. Sarris

## Abstract

The exocyst complex subunit protein Exo70B1 plays a crucial role in a variety of cell mechanisms including immune responses against pathogens. The calcium dependent kinase 5 (CPK5) of *Arapidopsis thaliana*, phosphorylates *At*Exo70B1 upon functional disruption. We previously reported that, the *Xanthomonas campestris* pv. *campestis* effector XopP, compromises Exo70B1 and bypasses the host’s hypersensitive response (HR), in a way that is still unclear.

Herein we designed an experimental approach based on biophysical, biochemical and molecular assays, based on structural and functional predictions, as well as, utilizing Aplhafold and DALI online servers respectively, in order to characterize the *in vivo Xcc*XopP function.

The interaction between *At*Exo70B1 and *Xcc*XopP is very stable in high temperatures, while the *At*Exo70B1 appeared to be phosphorylated at *Xcc*XopP expressing transgenic *Arabidopsis*. *Xcc*XopP reveals similarities with known mammalian kinases, and phosphorylates *At*Exo70B1 at Ser107, Ser111, Ser248, Thr309 and Thr364. Furthermore, *Xcc*XopP protects *At*Exo70B1 from AtCPK5 phosphorylation.

Together these findings show that, *Xcc*XopP is an effector, which not only functions as a novel serine/threonine kinase upon its host’s protein target *At*Exo70B1, but also protects the latter from the innate AtCPK5 phosphorylation, to bypass the host’s immune responses.

## Introduction

Plant pathogens of the *Xanthomonas* bacterial genus, originally include over 125 pathovars expanding in a 400 plant-host range (Timilsina *et al*., 2020). The pathovar “campestris” is the causal agent of black rot of cruciferous plants and together with the pathovars “raphani” (bacterial spot agent) and “incanae” (bacterial blight agent) form the three main pathovars, based on pathogenicity profiles (Vicente and Holub, 2013). The plant pathogen *Xanthomonas campestris* pv. *campestris* genome was the first published in 2002, revealing a number of mechanisms for host adaptation (White *et al*., 2009). Specifically, the type two secretion system (T2SS) contributes to host adaptation by secretion of cell wall degrading enzymes, while the type three, type four and type six secretion systems (T3SS, T4SS and T6SS, respectively), secrete effectors, with T3SS being the major type in pathogenicity (Tang *et al*., 2021). The number of T3SS effectors secreted by *Xanthomonas campestris* pv. *campestris* stands almost at 40 groups, with only XopP, XopAL1 and XopF1 in its core effectorome (Roux *et al*., 2015).

Effectors are injected by pathogens inside host plant cells, in order to overcome the host defense responses (Büttner and He, 2009). The first layer of defense is the recognition of pathogen or microorganism-associated molecular patterns (PAMPs or MAMPs), resulting in the PAMP-triggered immunity (PTI) (Jones and Dangl, 2006). Subsequently, host can recognize the pathogen effectors via specialized immunity receptors known as nucleotide binding leucine-rich repeat receptors (NB-LRR or NLR), which triggers the effector-triggered immunity (ETI) (Marchal *et al*., 2022; Mermigka and Sarris, 2019; Jones and Dangl, 2006). This two-layered system leads to a variety of responses, such as accumulation of oxygen species, callose deposition, mitogen activated protein kinase activation and pathogenesis related gene expression (Sertedakis *et al*., 2022; Yuan *et al*., 2021).

Tethering of secretory vehicles to the cell membrane is performed by the exocyst complex prior to SNARE-mediated fusion (Mei *et al*., 2018). The exocyst is a highly conserved, octameric complex, consisting by the SEC3, SEC5, SEC6, SEC8, SEC10, SEC15, Exo70 and Exo84 subunits (Mei and Guo, 2018). The Exo70 protein family appears to be the most divergent in plants, containing three subfamilies (Exo70.1; Exo70.2 and Exo70.3) and nine subgroups (Exo70A to Exo70I) (Zhao *et al*., 2019). The so far studied Exo70 groups have been proven to contribute in a variety of in cell procedures (Zhao *et al*., 2019).

It has been reported that the Exo70B1 and Exo70B2 paralogs are involved in a variety of cellular mechanisms such as stomatal opening and closure, autophagy and immune responses to different plant pathogens (Michalopoulou *et al*., 2022; Nishimura *et al*., 2017; Kulich *et al*., 2013; Hong *et al*., 2016). Particularly, the Exo70B1 subunit interacts with RIN4 and guides the exocyst complex to the plasma membrane (Sabol, Kulich and Žárský, 2017), which makes it a requirement for the early stages of immune signaling events, triggered by the bacterial flagellar peptide, flg22 (Wang *et al*., 2020). The importance of *At*EX070B1 in PTI defense responses makes this protein a hub-target for various pathogen effectors (Wang *et al*., 2019; Michalopoulou *et al*., 2022; Tsakiri *et al*., 2022). This could explain the guarding of *At*Exo70B1 by the NLR immunity receptor TN2 (Zhao *et al*., 2015). The interaction of TN2 and CPK5 (Calcium-dependent kinase 5) is responsible the hypersensitive response (HR) activation when the *At*Exo70B1 is compromised (Liu *et al*., 2017).

It is well established that effectors’ structural biology studies can be a revealing factor, changing the concept of creating novel strategies against pathogens (LeBlanc *et al*., 2021; Mak and Thurston, 2021; Kotsaridis, Tsakiri and Sarris, 2022). Although the classical techniques for protein structure identification can be difficult and, in some cases impossible, the rapid development of artificial intelligence software, like Alphafold, can be very accurate for protein structures’ prediction (Tunyasuvunakool, 2022).

Earlier this year, Michalopoulou et. *al*., reported that the *Xanthomonas campestris* pv. *campestris* type three effector XopP (here after *Xcc*XopP) to interact with *Arabidopsis thaliana* Exo70B1 protein (here after *At*Exo70B1) by their N-termini (Michalopoulou *et al*., 2022). This discovery was the result of a large interactions screening between the *Xcc*XopP and a number of NLR integrated decoys (IDs), including an Exo70-like ID (integrated decoy) of RGH2 NLR from *Hordeum vulgare* subsp. *vulgare* (Brabham *et al*., 2018). NLR IDs function as bait, luring T3SS effectors and triggering host’s immune responses, which is the reason of their high conservation (Daskalov *et al*., 2015; Sarris *et al*., 2016; Sandstrom *et al*., 2019). This interaction led the authors to identify a subcellular target of *Xcc*XopP which was the *At*Exo70B1 protein. The *Xcc*XopP-*At*Exo70B1 interaction, disrupts the exocyst subcomplex-ll formation, as long as the interaction of the two subcomplexes (Ahmed *et al*., 2018; Mei *et al*., 2018; Synek *et al*., 2021; Michalopoulou *et al*., 2022). Moreover, the secretion of several PTI related components that use the exocyst complex for their secretion/translocation, such as the pathogenesis-related protein-1A (PR1a) (Gu and Innes, 2012; Du *et al*., 2015; Michalopoulou *et al*., 2022), callose deposition (Du *et al*., 2015; Redditt *et al*., 2019; Michalopoulou *et al*., 2022) and the PRR immunity receptor FLS2 (Wang *et al*., 2020; Michalopoulou *et al*., 2022), was inhibited. The in-host mechanism that governs such a successful effector, disrupting PTI responses without triggering TN2-dependent immune responses is something that is very important to be investigated.

In the present study, using biophysical, biochemical and molecular approaches, as well as, structural and functional predictions, we provide evidence that *Xcc*XopP not only interferes with the phosphorylation of *At*Exo70B1 by AtCPK5, by a very stable interaction that inhibits HR induction, but also phophorylates *At*Exo70B1 by itself, revealing new insights in *Xcc*XopP functions in the host cell.

## Material and methods

### Structure prediction analysis

The 3D structures of all the studied proteins in this study were predicted using the artificial intelligence (AI) program AlphaFold that it is designed as a deep learning system for proteins structure prediction. (Jumper *et al*., 2021; Varadi *et al*., 2022). Subsequently, the predicted structure was compared with those in the Protein Data Bank via the Dali online server (Holm, 2020). The visualization of the protein structures was performed by PyMOL (‘The PyMOL Molecular Graphics System, Version 2.0 Schrödinger, LLC.’).

### Plant materials and growth conditions

*Arabidopsis thaliana* ecotype Columbia-0 (Col-0) and *Xcc*XopP transgenics (Michalopoulou *et al*., 2022) were used in our study for NanoLC/MS-MS analysis. Arabidopsis seeds were stratified at 4°C in water for 3 days before sowing in soil and then were grown in greenhouse conditions.

### Confocal microscopy

All samples were imaged with a 40x water objective. The confocal images were acquired in a SP8 Leica confocal microscopy and processed with ImageJ. mCherry was excited using a 561 nm laser and emission was collected at 593-628 nm. Chlorophyll was detected between 653-676 nm and excited at 561 nm.

### NanoLC-MS/MS Analysis

The nanoLC-MS/MS analysis was performed on an EASY-NanoLC II system (Thermo Scientific) coupled with an LTQ-Orbitrap XL ETD (Thermo Scientific) through an ESI ion source (Thermo Scientific). Data were acquired with Xcalibur software (LTQ Tune 2.5.5 sp1, Thermo Scientific). Prior to the analysis, the mass spectrometer was calibrated with a standard ESI positive ion calibration solution of caffeine (Sigma), L-methionyl-arginyol-phenylalanylalanine acetate H_2_O (MRFA, Research Plus, Barnegat, NJ), and perfluoroalkyl triazine (Ultramark 1621, Alfa Aesar, Ward Hill, MA). Samples were reconstituted in 0.5% formic acid, and the tryptic peptide mixture was separated on a reversed-phase column (Reprosil Pur C18 AQ, particle size = 3 μm, pore size = 120 Å (Dr. Maisch, AnaLab, Athens, Greece), fused silica emitters 100 mm long with a 75 μm internal diameter (New Objective) packed in-house using a pressurized (35 to 40 bars of helium) packing bomb. The nanoLC flow rate was 300 nl*min^-1^. The LC mobile phase consisted of 0.5% formic acid in water (A) and 0.5% formic acid in acetonitrile (B). A multi-step gradient was employed, from 5% to 30% B for 120 min, to 90% B for 10 min. After the gradient had been held at 90% B for 5 min, the mobile phase was re-equilibrated at initial gradient conditions. The MS was operated with a spray voltage of 2,300 V, a capillary voltage of 35 V, a tube lens voltage of 140 V, and a capillary temperature of 180 °C. A survey scan was acquired in the range of *m/z* 400–1,800 with an AGC MS target value of 10^6^ (resolving power of 60,000 at *m/z* 400). The 10 most intense precursor ions from each MS scan were subjected to collision-induced dissociation in the ion trap.

### Data Analysis of MS/MS-derived Data

The MS raw data were loaded in Proteome Discoverer 2.2.0 (Thermo Scientific) and run using Mascot 2.3.02 (Matrix Science, London, UK) and Sequest HT (Thermo Scientific) search algorithms against the *At*EXO70B1 (NM_125229.4) sequence. A list of common contaminants was included in the database. For protein identification, the following search parameters were used: precursor error tolerance = 10 ppm, fragment ion tolerance = 0.8 Da, trypsin full specificity, maximum number of missed cleavages = 3, and cysteine alkylation as a fixed modification, Oxidation of Methionine, Acetylation of N-term, Phosphorylation (The node ptmRS was used) of Serine, Threonine and Tyrosine as Variable Modifications. To calculate the protein false discovery rate, a decoy database search was performed simultaneously with strict criteria set to 0.01 and relaxed criteria to 0.05.

### Protein production and purification

**XccX*opP, *XccX*opP* fragment (100-733 aa, *Xcc*XopP_100_) and **Xcc*XopP* fragment (1-420 aa, *Xcc*XopP_1-420_) genes were cloned and inserted into the pET26b (ColE1 plasmids) vector carrying a C-terminal 6-His tag and transformed into the *E.coli* strain BL21. A sufficient amount of soluble protein was obtained after induction using the following conditions: Cells were grown in LB medium containing 30 μg/ml kanamycin until an OD_600_ value of 0.6-0.8 was reached. The culture was induced with 0.5 mM IPTG and left to grow at 20°C overnight. The cell paste was re-suspended in 100 ml lysis buffer containing 25 mM Tris pH 8.0, 300 mM NaCl, 5 mM imidazole and 15 mM β-mercaptoethanol, and homogenized. After adding protease inhibitors (20 μg/ml leupeptin, 1 mM PMSF and 150 μg/ml benzamidine), the cells were disrupted with sonication on ice, 10 sonication cycles of 30 s each, with cooling intervals of 30 s. The precipitate was subsequently removed by centrifugation at 12,000 rpm at 4°C for 45 min. Purification was carried out using His-tag affinity chromatography at 4 °C with an 8 ml Ni-NTA Qiagen column pre-equilibrated in lysis buffer and initially washed stepwise with 10, 20 and 30 mM imidazole. With a subsequent increase in imidazole concentration the protein started eluting at 100 mM imidazole. Fractions containing the protein were dialyzed against the storage buffer containing 25 mM Tris pH 8.0, 100 mM NaCl and 10 mM β-mercaptoethanol. In case of *At*Exo70B1, the full length protein was cloned and inserted into the LIC 1.10 vector carrying an N-terminal 6-His-SUMO3 tag, and transformed into the *E.coli* strain Rosetta™ (DE3). The same protocol was followed and the fractions containing the protein were dialyzed against the storage buffer (25 mM Tris pH 8.0, 100 mM NaCl, 10 mM β-mercaptoethanol and 0.5 nM SENP-2 protease). Subsequently, a reverse purification protocol was carried out using His-tag affinity chromatography. The elution fractions containing the proteins, were pooled and concentrated to a final volume of 2 ml. Size exclusion chromatography was performed in 20 °C ÄKTA purifier system (Amersham) and a prepacked Hi-Prep 16/60 Sephacryl S-200 high-resolution column (GE Healthcare). Flow rate was 0.5 ml/min, and elution was monitored at 280 nm. Using a 2 ml-loop the proteins were loaded, and the 2 ml fractions were collected and analyzed via a 10% SDS-polyacrylamide gel.

### Protein extraction from plant leaves

WT and *Xcc*XopP transgenic Arabidopsis leaves were excised from the plants and ground in liquid nitrogen. Proteins were extracted in GTEN buffer (10% glycerol, 25 mM Tris-Cl pH 7.5, 1 mM EDTA, 150 mM NaCl) supplemented with fresh 5 mM DTT, protease inhibitor cocktail (Sigma, P9599) and 0.15% v/v NP-40. After 10 min centrifugation at 4°C at 14,000 x g, the supernatant was collected in a clean microcentrifuge tube.

### SEC-MALS Analysis

After purification, SEC-MALS—the combination of size-exclusion chromatography with multi-angle light scattering, was used to monitor the oligomerization states of *Xcc*XopP. For all the proteins, the analysis was performed as follows: 100 μL from the samples were loaded onto Superdex 200 columns (GE Healthcare) connected to a high-performance liquid chromatography (HPLC) system (Shimadzu) operating with the LC solution software equipped with a solvent delivery module (Shimadzu; LC-20AD), a UV/VIS photodiode array detector (Shimadzu; SPD-M20A) measuring at 280 nm, a differential refractive index detector (Shimadzu; RID-10A), and a system controller (Shimadzu; CBM-20A) and coupled to online mass detection by an advanced 8 angles MALS detector (Wyatt; Dawn 8+) with an integrated Wyatt QELS Dynamic Light Scattering (DLS) module. Data were analyzed with the Astra software (ASTRA 6.1.2.84).

### Field-Emission Scanning Electron Microscopy (FE-SEM)

The *Xcc*XopP_100_ fragment was purified by His-tag affinity chromatography. Fractions containing the protein were dialyzed against the storage buffer in the absence of β-mercaptoethanol overnight. The gel-like form that came up, was deposited on a covered glass and air dried overnight. The sample was then covered with 10 nm of Au/Pd sputtering and observed using a JEOL JEM-2100 transmission electron microscope at 20 kV.

### Transmission Electron Microscopy (TEM)

5 μl sample from *Xcc*XopP_100_ was deposited onto a formvar/carbon-coated electron microscopy grid for 2 min. After removing the excess with filter paper, 2% w/v uranyl acetate was used in order the sample to be stained, for 2 min. The observations were conducted with a JEOL JEM-2100 transmission electron microscope at 80 kV.

### Circular Dichroism Measurements

CD spectra for *Xcc*XopP, *At*Exo70B1 and *Xcc*XopP-*At*EXO70B1 complex were also obtained using a J-810 CD spectropolarimeter (Jasco Inc.) with quartz cuvettes of 1-mm path length and a protein concentration of 0.5 mg/mL using a 0.1-mm demountable quartz cuvette. Thermal denaturation was monitored by the change of the CD signal at 222 nm for 10–90 °C and a waiting time of 2 s for stabilization. The Spectra Manager program (Jasco Corp.) was used for buffer subtraction and unit conversions to MRE.

### Nano Differential Scanning Fluorimetry (DSF)

NanoDSF was performed using Prometheus NT.48 equipped with back reflection mode (NanoTemper Technologies, München, Germany). *Xcc*XopP, *At*Exo70B1 and their complex were loaded in nanoDSF grade standard capillaries (NanoTemper Technologies GmbH, München, Germany) and exposed at thermal stress from 20 ^⍰^C to 95 ^⍰^C by thermal ramping rate of 1 ^a^C/min. Fluorescence emission from tryptophan after UV excitation at 280 nm was collected at 330 nm and 350 nm with dual-UV detector.

#### *In vitro* Protein Kinase Assay

For the *in vitro* AtCPK5 and *Xcc*XopP phosphorylation assay, the reaction was performed according to Liu *et al* with modifications (Liu *et al*., 2017). Reactions were prepared, with a 30 min incubation of *Xcc*XopP and *At*Exo70B1 before adding *At*CPK5 to sample. The samples then were tested for phosphorylation sites with NanoLC-MS/MS Analysis.

## Results

### The XccXopP lacking the first 100 amino acids self-assembles into fibrils

In order to investigate the large unfolded N-terminal region of *Xcc*XopP, a fragmented version of *Xcc*XopP containing the region of 100 aa to 733 aa (hereafter *Xcc*XopP_100_), was expressed and purified. Size exclusion chromatography and SEC-MALS analysis revealed that the truncated *Xcc*XopP_100_ was able to form different molecular size multimers (Fig. 1A, B). Another interesting fact is that, during the concentration stage of *Xcc*XopP_100_, a gel-like product was observed in the protein sample. Transmission electron microscopy (Fig. 1C) and scanning electron microscopy (Fig. 1D) was performed to the gel-like protein product that was obtained after the sample’s multimarization. Our analysis revealed that the truncated *Xcc*XopP_100_ has the ability to self-assemble into fibrils, suggesting that the N-terminal unfolded region of *Xcc*XopP probably protects the effector from oligomerization.

**Figure 1.**
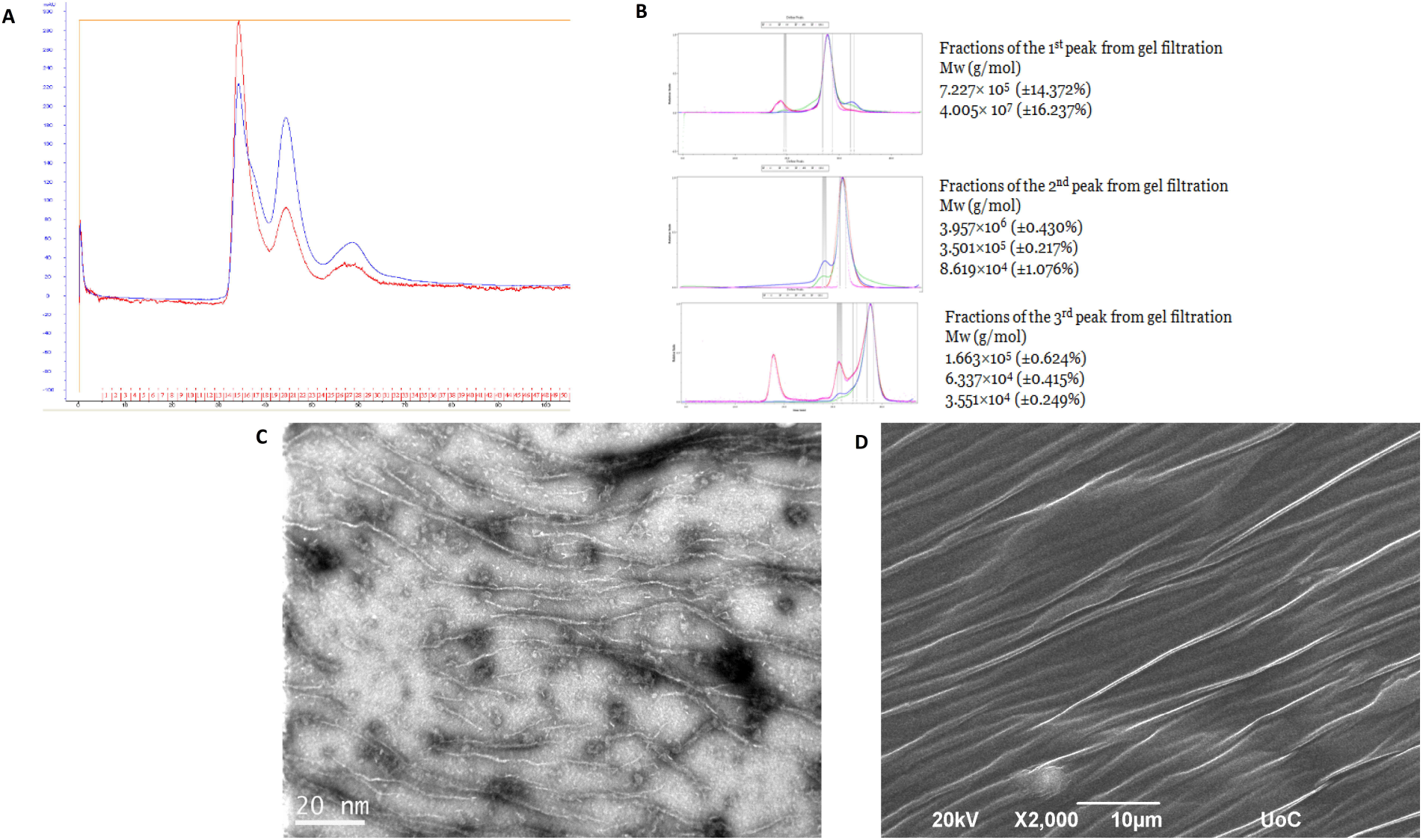
*Xcc*XopP_100_ fragment self-assembles into fibrils: **A.** *Xcc*XopP_100_ size exclusion chromatography diagram, **B**. SEC-MALS on the different elution peaks (earlier elution volumes from top to bottom), **C**. Transmission electron microscopy image of the concentrated *Xcc*XopP_100_ protein sample, **D**. Scanning electron microscopy of *Xcc*XopP_100_ gel-like product.

### Structural prediction of *At*Exo70B1 and *Xcc*XopP

The *At*Exo70B1 protein structure was predicted using the AI program Alphafold (Fig. 2A). The tertiary structure appears as an elongated molecule containing only *α*-helices. Subsequently, the predicted structure of *At*Exo70B1 was superimposed to the known structure of *At*Exo70A1 (Fig. 2B and 2C). The N-termini of the two proteins appear to be differentiated. The first 93 residues of *At*Exo70A1 form a single α-helix, where the first 54 residues of *At*Exo70B1 form 2 α-helices connected to the rest of the molecule with a short loop. The superimposition between these protein molecules resulted in a root mean square deviation (RMSD) value of 1.8, with a major variation to be the N-termini and at a loop region that seemed to be larger in the case of *At*Exo70B1 (Fig. 2C).

**Figure 2.**
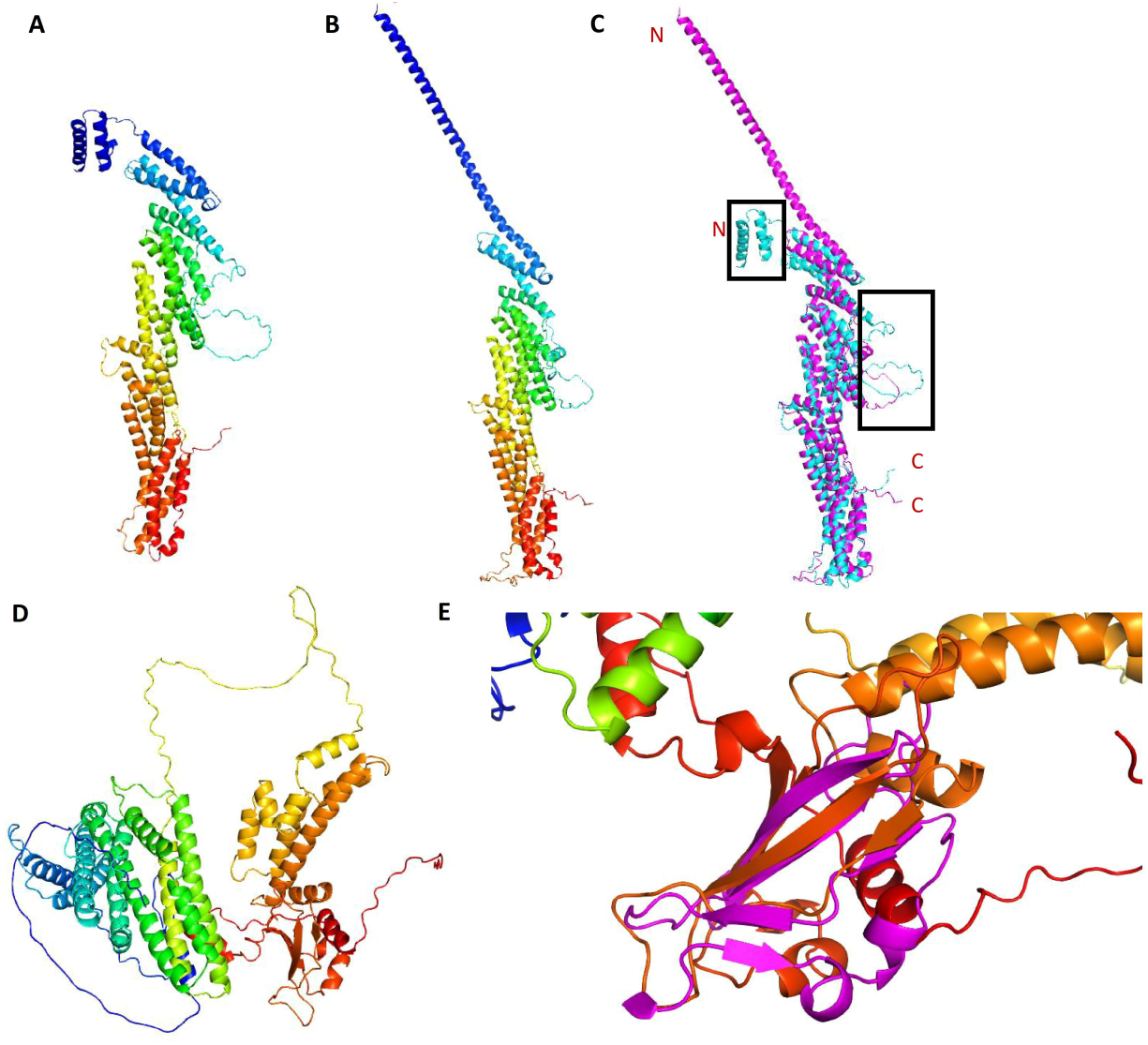
3D Structure prediction: **A.** *At*Exo70B1 (N-terminus blue, C-terminus red), **B.** *At*Exo70A1 (N-terminus blue, **C.** Superimposition between *At*EXO70B1 (cyan) and *At*EOX70A1 (magenta). Frames indicate the loops and the N-terminus variations between the two proteins, **D**. *Xcc*XopP (N-terminus blue, C-terminus red), **E**. Superimposition between *Xcc*XopP (multicolor) and mammalian mTOR KD domain (magenta).

Additionally, the structure prediction of the Exo70-like integrated decoy of the NLR immune receptor *Hv*RGH2, was predicted (Fig. S1A) and superimposed to the *At*Exo70B1 (Fig. S1B). This resulted in high structural similarity between the Exo70-like ID and Exo70B1, with the differences to appear to their N-termini and to the loop region, similarly to the *At*Exo70B1 and *At*Exo70A1 comparison.

On the other hand, the 3D structure prediction of *Xcc*XopP revealed a two-domain molecule connected with a large loop (Fig. 2D). The N-terminus of this effector contains a large unfolded region and ten α-helices (Fig. 2D). The C-terminus consists of a number of short α-helices and loops, but also a four-stranded β-sheet interrupted by two small α-helices (Fig. 2D). The predicted *Xcc*XopP structure was compared to those deposited to Protein Data Bank using the DALI online server. The first hits of this search were the mammalian targets of rapamycin, mTOR (PDB: 7PEA) (Yang *et al*., 2013). The latter is a serine/threonine phosphoinositide-3 kinase. The superimposition of *Xcc*XopP C-terminus with mTOR kinase domain (KD) revealed a value of RMSD=2,763 (Fig. 2E).

Recently, it was reported that the *Xanthomonas oryzae* pv. *oryzae* XopP effector (*Xoo*XopP) interacts with the PUB44 E3 ligase from *Oryza sativa* and inhibit its ligase function (Ishikawa *et al*., 2014). In order to investigate the structural similarity of *Xcc*XopP with homologs of other bacterial pathogens, the tertiary structures of: *Xanthomonas oryzae* pv. *oryzae* (Xoo) XopP-1 and XopP-2 (Lee *et al*., 2005) and the *Xanthomonas oryzicola* pv. *oryzicola* (Xor) XopP-1 and XopP-2 effectors (Bogdanove *et al*., 2011), as well as, the homologs from *Ralstonia solanacearum (Rs)* HLK3 (Salanoubat *et al*., 2002) and the *Acidovorax citrulli* (Ac) XopP (Baxevanis, 2000) effectors, were predicted and superimposed with *Xcc*XopP (Fig. S2). This resulted to a conserved β-sheet C-terminal domain but a high divergent N-termini, regarding the number of α-helices and loops length.

### *Xcc*XopP and *At*Exo70B1 form a complex with high thermal stability

For the further investigation of the dynamics of the *Xcc*XopP-*At*Exo70B1 complex, CD and nanoDSF were used for both proteins individually, as well as, for their complex. The melting temperatures (TM) of *Xcc*XopP and *At*Exo70B1 are approximately 40 °C and 50 °C, respectively. According to nanoDSF, the *Xcc*XopP-*At*Exo70B1 complex reveals two distinct TMs, almost identical to those of *Xcc*XopP and *At*Exo70B1 proteins (Fig. 3E). The CD spectra revealed the characteristic minima at 208 and 222 nm and a positive band at 195 nm, which is the common signature for α-helical proteins (Fig. 3A-C). Furthermore, the θ222/θ208 offers additional information about the α-helicity of the two proteins. In case of *At*Exo70B1, the value of the ratio is bigger than 1 (Fig. 3A), which suggests a conformation similar to coiled-coil. Regarding the *Xcc*XopP, the ratio is below 1, resulting in isolated α-helices. Another interesting point is the existence of two dichroic points for the complex (Fig. 3C), in comparison to solitary *Xcc*XopP and *At*Exo70B1 proteins. Taking in consideration the multiple different states that appear in the case of the complex due to the two dichroic points, along with its thermal stability revealed by the CD spectra presentation in different temperatures (Fig. 3D), *Xcc*XopP-*At*Exo70B1 complex seems to unfold and refold, as the temperature increases, suggesting a very strong interaction.

**Figure 3.**
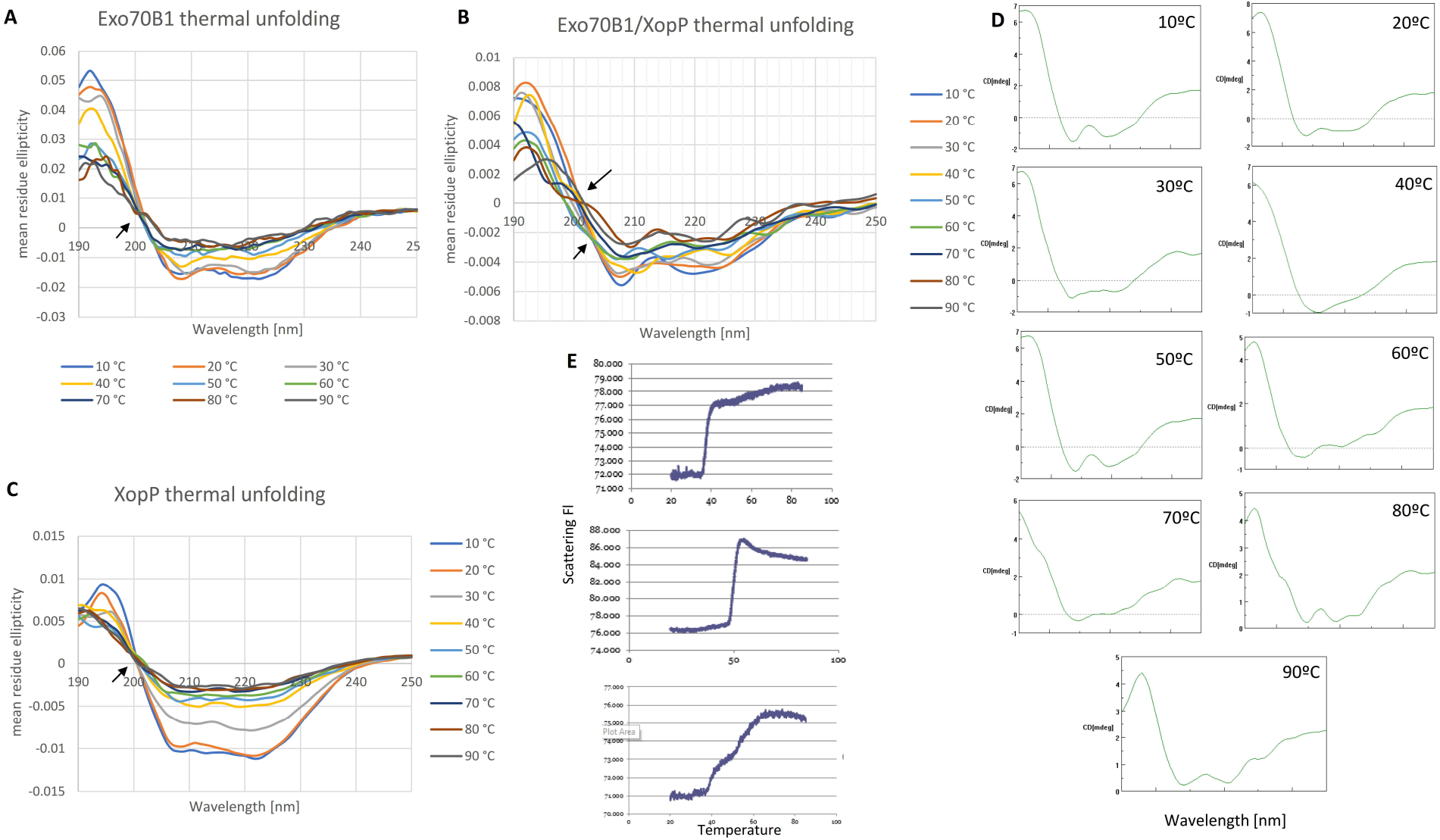
*Xcc*XopP and *At*Exo70B1 forms a complex with high thermal stability: **A.** *At*Exo70B1 CD spectra, **B.** *Xcc*XopP-*At*Exo70B1 complex CD spectra, **C.** *Xcc*XopP CD spectra, D. *Xcc*XopP-*At*Exo70B1 complex CD spectra of 10-90ºC temperature range, **E.** nanoDSF to *Xcc*XopP, *At*Exo70B1 and *Xcc*XopP-*At*Exo70B1 complex (top to bottom). Arrows indicate isodichroic points.

### *At*Exo70B1 appears phosphorylated in *Xcc*XopP transgenic plants

In order to investigate the potential effect that *Xcc*XopP could have on *At*Exo70B1, total protein extracts were obtained from both w.t., as well as, *Xcc*XopP expressing transgenic Arabidopsis plants (Michalopoulou *et al*., 2022). The *Xcc*XopP expression in transgenic Arabidopsis was confirmed via confocal microscopy, since the protein was tagged with an mCherry tag (Fig. 4A). The comparison analysis of nanoLC/MS-MS data between the w.t. and the transgenic plants revealed a greater number of phosphorylated amino acid residues for *At*Exo70B1 in transgenic plants, compared to wild type plants (Fig. 4B). We purified *Xcc*XopP and *At*Exo70B1 for *in vitro* assays, in order to investigate whether these phosphorylated sites are the result of *Xcc*XopP activity or of the native *Arabidospis thaliana* kinases.

**Figure 4.**
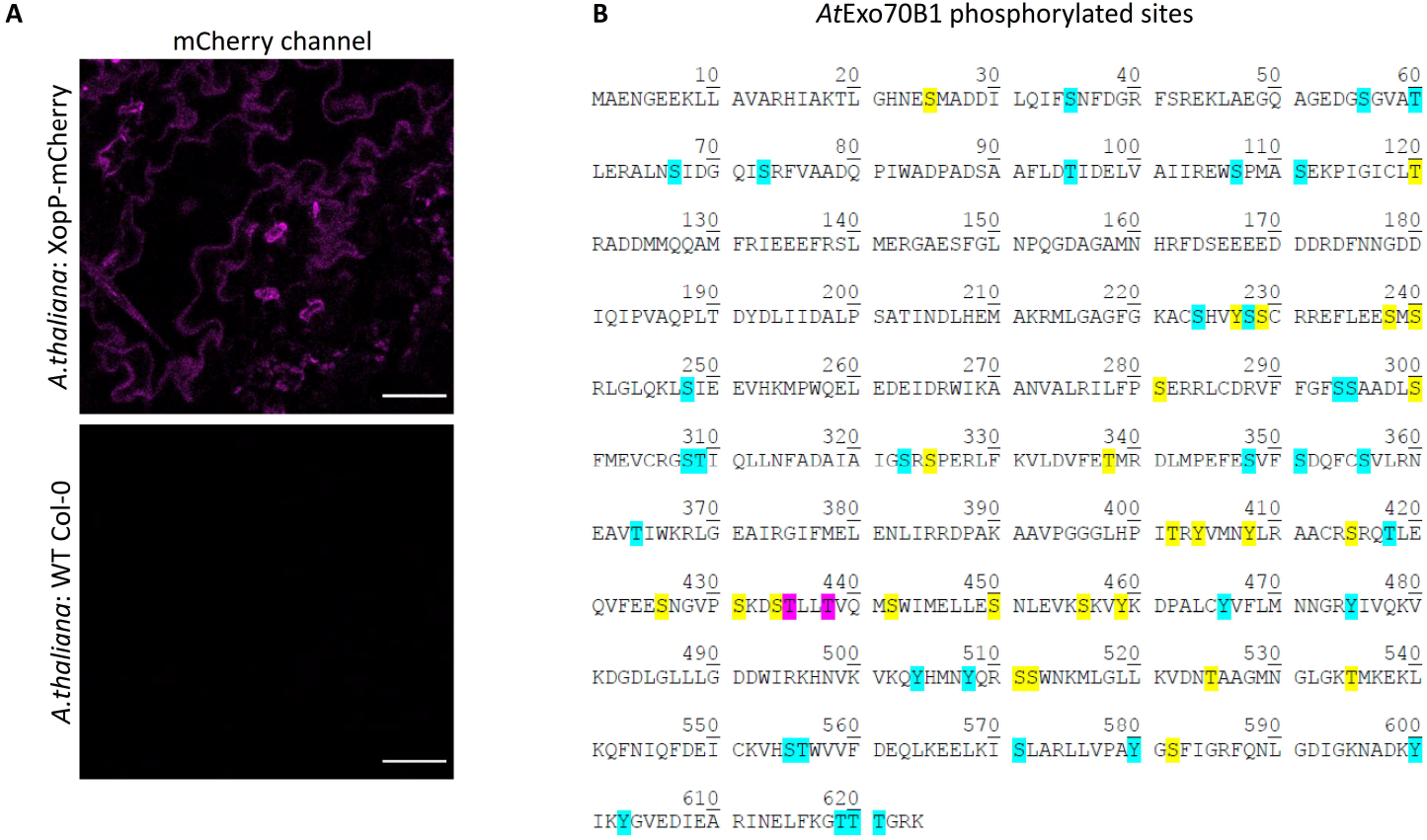
Phosphorylated residues of *At*Exo70B1 from WT and *Xcc*XopP transgenic *Arabidopsis thaliana* plants. **A**. Expression of XopP transgene in *Arabidopsis thaliana* transgenic plants. mCherry channel is shown with magenta color. Bars 50 Um, **B.** Phosphorylated sites highlighted in *At*Exo70B1 as cyan (transgenic plants), yellow (both in wild type and transgenic plants) and magenta (only in wild type).

### *Xcc*XopP phosphorylates *in vitro* the *At*Exo70B1

*In vitro* kinase assays were performed to elucidate *Xcc*XopP function to its host’s target *At*Exo70B1. NanoLC MS/MS data analysis revealed that *Xcc*XopP is capable to phosphorylate Exo70B1. The related phosphorylated aa residues of Exo70B1 were the: Ser107, Ser111, Ser248, Ser308, Thr309 and Thr364 (Fig. 5A-B) (Fig. S3) (Table S1). These findings come in agreement with our previously *in-vivo* experiments. These residues appeared phosphorylated only in transgenic plants. The truncated version of *Xcc*XopP_1-420_, which contains only the N-terminus of the effector protein, was not able to phosphorylate *At*Exo70B1 to any residue (Fig. S4), which suggests that the kinase activity exists in the C-terminus of *Xcc*XopP. Lastly, it is very interesting to mention that the residues that are phosphorylated appear to be in the domain of *At*Exo70B1 that interacts directly with *Xcc*XopP (Fig 5C), as it was predicted with Alphafold.

**Figure 5.**
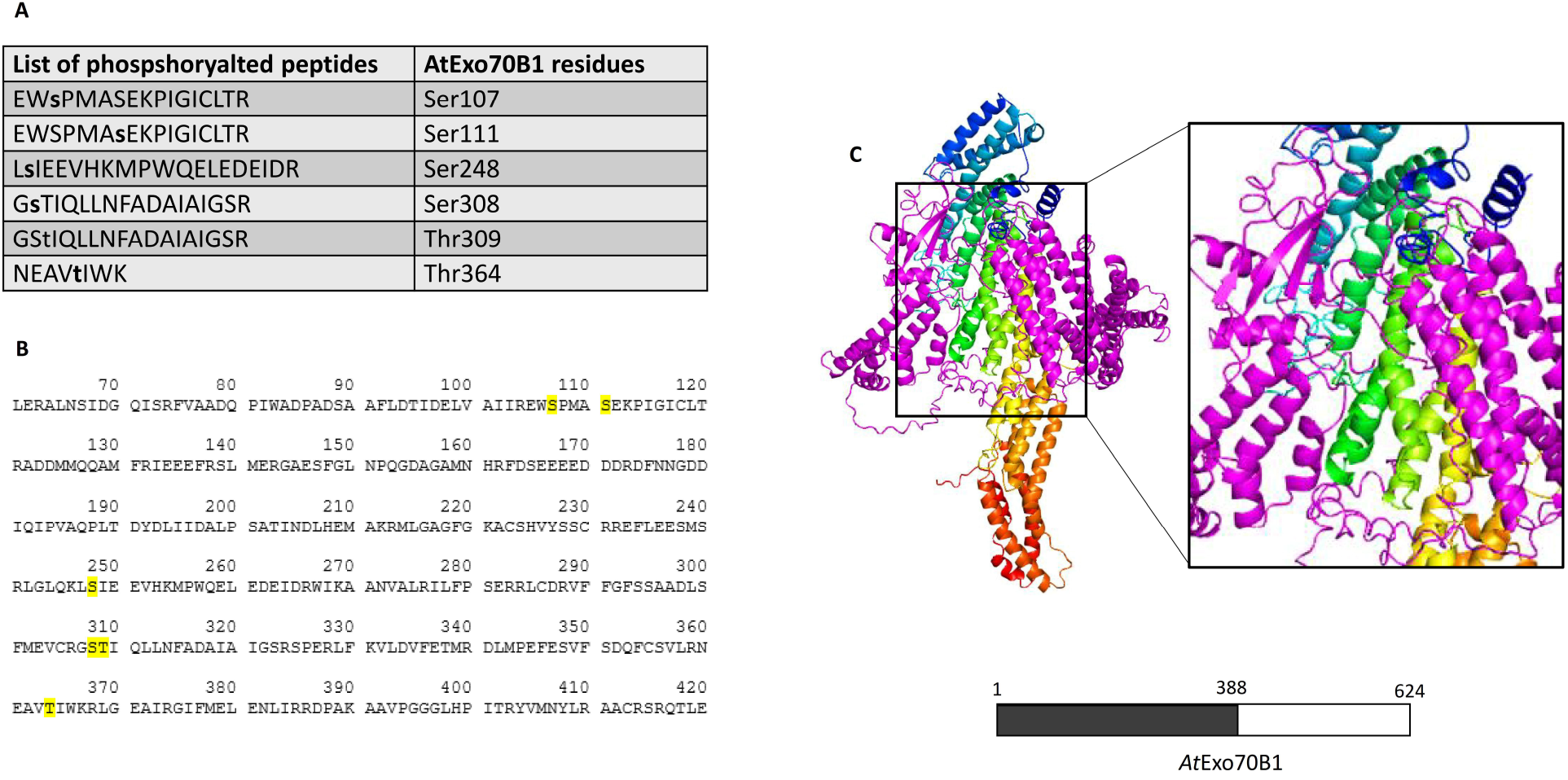
*Xcc*XopP phosphorylates *At*Exo70B1 *in vitro*. **A**. List of peptides containing phosphorylated residues, **B**. position of the phosphorylated residues upon *At*Exo70B1 amino sequence, **C**. Structure prediction of *Xcc*XopP (magenta)-*At*Exo70B1(multicolor) protein complex. The region of *At*Exo70B1 that interacts with *Xcc*XopP is indicated by grey color.

### *Xcc*XopP protects *At*Exo70B1 from AtCPK5 phosphorylation

It has already been proven by Liu et al, that *At*CPK5 phosphorylates *At*EXO70B1 when it is compromised (Liu *et al*., 2017). The MS/MS derived data revealed that, AtCPK5 phosphorylates the residues: 19Thr, 25Ser, 67Ser, 364Thr, 582Ser residues of *At*Exo70B1 in consistency and with high confidence (Fig. 6A) (Table S2). The same results were obtained when *Xcc*XopP, Exo70B1 and AtCPK5 were added at the same time in the protein mixture. However, the related residues were not phosphorylated when a 30 minutes pre-incubation of *At*Exo70B1 and *Xcc*XopP were performed prior to the AtCPK5 addition (Fig. 6A). Interestingly, the N-terminus of the effector seems to have the same effect on *At*Exo70B1 (Table S2). This suggests that *Xcc*XopP disrupts the proper interaction of AtCPK5 with *At*Exo70B1 and its phosphorylation. These results come to an agreement with the protein complex of *At*Exo70B1-AtCPK5 predicted structure for the phosphorylated of N-terminal residues (Fig. 6B).

**Figure 6.**
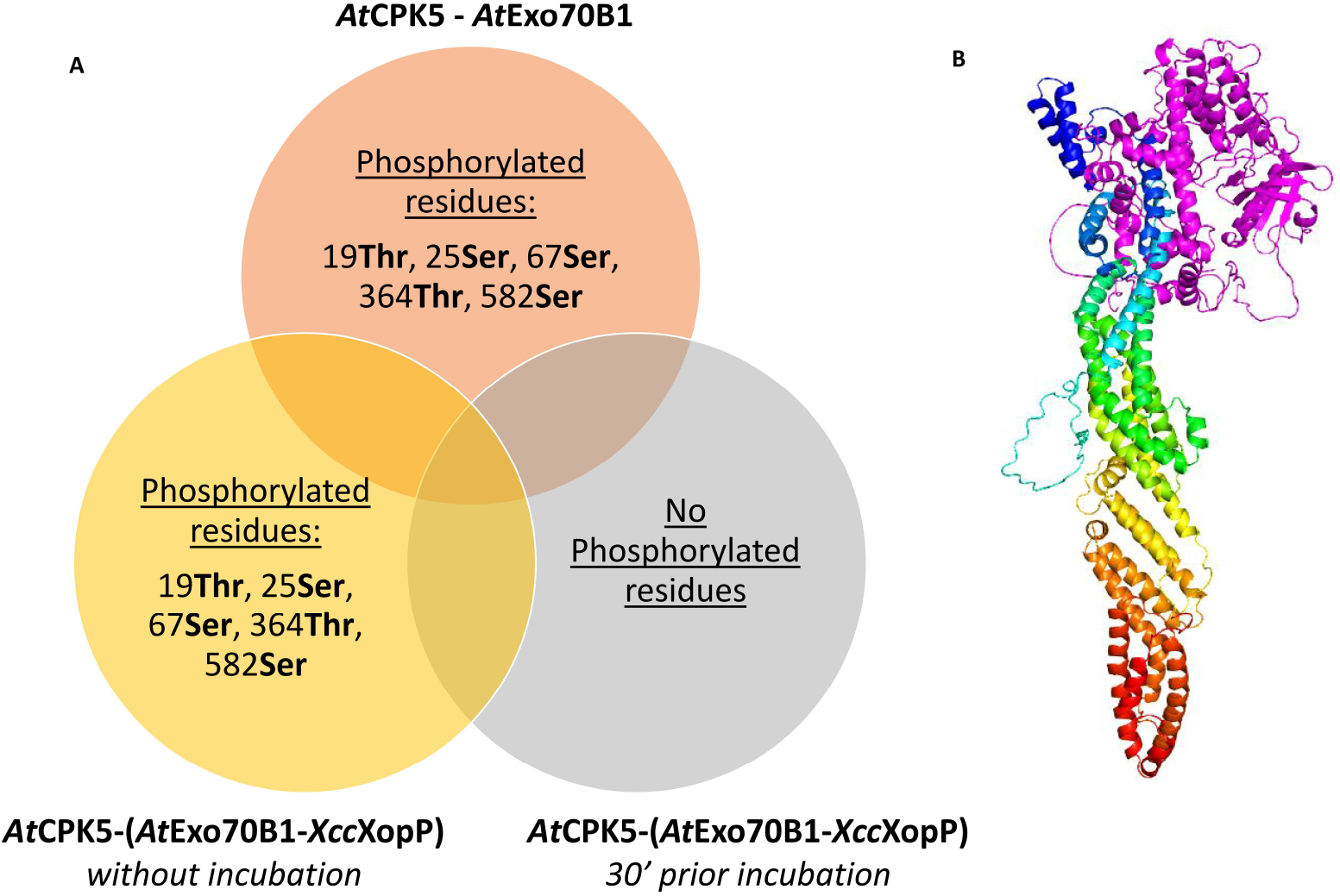
*Xcc*XopP protects *At*Exo70B1 from AtCPK5 phosphorylation in vitro. **A.** Common high confident phosphorylated peptides from *in vitro kinase* assays: The *At*Exo70B1 phosphorylated residues by AtCPK5 (orange circle), from *in vitro* kinas assay, are identical when we perform the assay with AtCPK5-*At*Exo70B1-*Xcc*XopP without a prior incubation (yellow circle). No phosphorylated residues were observed, when *At*Exo70B1-*Xcc*XopP were incubated in the protein sample for 30 minutes, **B**. Structure prediction of AtCPK5 (*magenta*)-*At*Exo70B1 (multicolor) protein complex.

## Discussion

The exocyst complex is a conserved octameric protein complex that regulates the tethering of secretion vesicles to the cell membrane (Mei and Guo, 2019). Naturally, the components of such an important cellular mechanism, would make the exocyst complex components potential virulence targets. Exo70B1 is one of them and belongs to the family of Exo70 proteins (Stegmann *et al*., 2014). Several studies have proven that the Exo70 protein groups are involved in a variety of processes inside the host cell (Zhao *et al*., 2019; Žárský *et al*., 2020). Many pathogens use protein effectors, which reveal a variety of functions. These effectors are injected into the host cells helping pathogens to bypass host defense mechanisms (Büttner and Bonas, 2003; Kotsaridis, Tsakiri and Sarris, 2022; Sertedakis et al., 2022).

The previously reported interaction between *Xcc*XopP and *At*Exo70B1 was capable to disrupt the subcomplex-II formation and moreover the exocyst complex final assembling (Ahmed *et al*., 2018; Mei *et al*., 2018; Synek *et al*., 2021; Michalopoulou *et al*., 2022). This exocyst functional disruption, leads to downregulation of a variety of PTI related responses, such as: callose deposition, FLS2 PRR translocation to the PM and PR1 protein secretion (Gu and Innes, 2012; Du *et al*., 2015; Redditt *et al*., 2019; Wang *et al*., 2020). Moreover, the lack of interaction ability of the exocyst subunits *At*Exo70B1 and *At*Exo84B, lead to their reduced protein levels. Lastly, *Xcc*XopP that can modify host’ PTI responses and simultaneously not triggering TN2-dependent immune response (Michalopoulou *et al*., 2022).

According to the above, we investigate further the biophysical and biochemical properties of the *Xcc*XopP-*At*Exo70B1 interaction, as long as the function of *Xcc*XopP upon its host’s protein target. Firstly, we utilized bioinformatics tools/software to predict the tertiary structure of the two proteins. The protein structure of *At*Exo70B1 in comparison of *At*Exo70A1, showed high homology with the main divergent to be localized at their N-termini of the proteins.

Regarding the T3S effector *Xcc*XopP, the structural prediction reveals that it harbors a very long N-terminal unfolded region (Fig. 2D). Furthermore, the comparison between the *Xcc*XopP and its homologs *Xoo*XopP1, *Xoo*XopP2, *Xor*XopP1, *Xor*XopP2, *Rs*HLK3 and *Ac*XopP resulted to a highly abundant in α-helices N-termini, a very larger loop separating the N-terminus from the C-terminus, which consists of a number of α-helices and a conserved β-sheet domain. The C-terminus of *Xcc*XopP resulted to have high structure homology to the mammalian mTOR protein, a serine/threonine kinase (Yang et al., 2013). During their life, plants need to regulate their cell mechanisms such as development, reproduction, immunity in response, etc. Protein phosphorylation is the most abundant post-translation modification that has a very crucial role in a number of biological procedures and its catalyzed by specific enzymes, which called kinases (Zulawski and Schulze, 2015) and has the ability to transfer phosphoryl groups from ATP molecules to their targeted substrates residues Ser, Thr and Tyr (van Wijk *et al*., 2014). Moreover 940 kinases are encoded from *Arabidopsis thaliana* genome (Zulawski *et al*., 2014). The high regulatory importance of such enzymes, lead to a variety of T3SS effectors to adopt such functions in order to interfere with their hosts developmental or immunity mechanisms.

Cyclical dichroism is a structural method that uses polarized light, measuring the differential absorption of left- and right-handed light, and it consists a standard method to study the biophysical properties not only for single protein molecules, but also for protein-protein interactions (Whitmore and Wallace, 2008). According to the above, the CD spectra minima showed that the two proteins are consisted of α-helices, forming a structure similar to coiled-coil for *At*Exo70B1. In the case of *Xcc*XopP, the CD spectra minima revealed that it consists of isolated α-helices, which is in contrast with the structure protein prediction. We believe that this is the result of the high number of α-helices that appear in the protein molecule. Subsequently, the complex seems to keep its formation, even at 90°C. The existence of a isodichroic point in CD spectra indicates a local two-state population for a protein sample (Holtzer and Holtzer, 1992). The appearance of two dichroic points in combination with the complex stability in high temperatures indicates that, the complex transits between two states (folding-unfolding) and keeps its formation as temperature increases.

Based on the difference to the sites of phosphorylated residues between *Arabidopsis thaliana* wild type plants and Arabidopsis *Xcc*XopP transgenic plants, in order to clarify whether these phosphorylated sites are the result of the *Xcc*XopP activity or of the large number of native kinases, the two proteins were expressed and purified for *in vitro* assays. The gel-like protein product, which appeared during the purification of the truncated *Xcc*XopP_100_ protein, suggested that the long unfolded N-terminal region protects the effector from aggregation.

The function of *Xcc*XopP upon *At*Exo70B1 was researched by *in vitro* kinase assays. Firstly, the *At*Exo70B1 Ser107, Ser111, Ser248, Ser308, Thr309 and Thr364 residues were phosphorylated by *Xcc*XopP, proving that the latter acts as a kinase on *At*Exo70B1. This was not the case for the truncated *Xcc*XopP_1-420_, which lacks the β-sheet C-terminal domain, proving that the effector C-terminus is responsible for the kinase activity. Taking in consideration our previous results, this could be the reason of *At*Exo70B1 downregulation in presence of *Xcc*XopP (Michalopoulou *et al*., 2022). Moreover, the phosphorylated sites of *At*Exo70B1 by *At*CPK5 (19Thr, 25Ser, 67Ser, 364Thr, 582Ser) were not observed, when *At*Exo70B1 and *Xcc*XopP were incubated prior of the kinase assay with AtCPK5. That suggests that the effector has the ability to disrupt the phosphorylation of *At*Exo70B1 by AtCPK5 and therefore not triggering the host’s immune responses (Michalopoulou *et al*., 2022).

Overall, using a standard experimental pipeline (protein structure prediction - comparison with online databases - *in vivo* or *in vitro* assays) we prove that, the bacterial pathogen *Xanthomonas campestris* pv. *campestris* utilizes the T3SS effector *Xcc*XopP, a novel bacterial kinase, to bypass its host *Arabidopsis thaliana* defense response, while it simultaneously disrupts exocyst subunit *At*Exo70B1 function. The proposed mechanism, as it was created with Alphafold, is represented in Figure 7.

**Figure 7.**
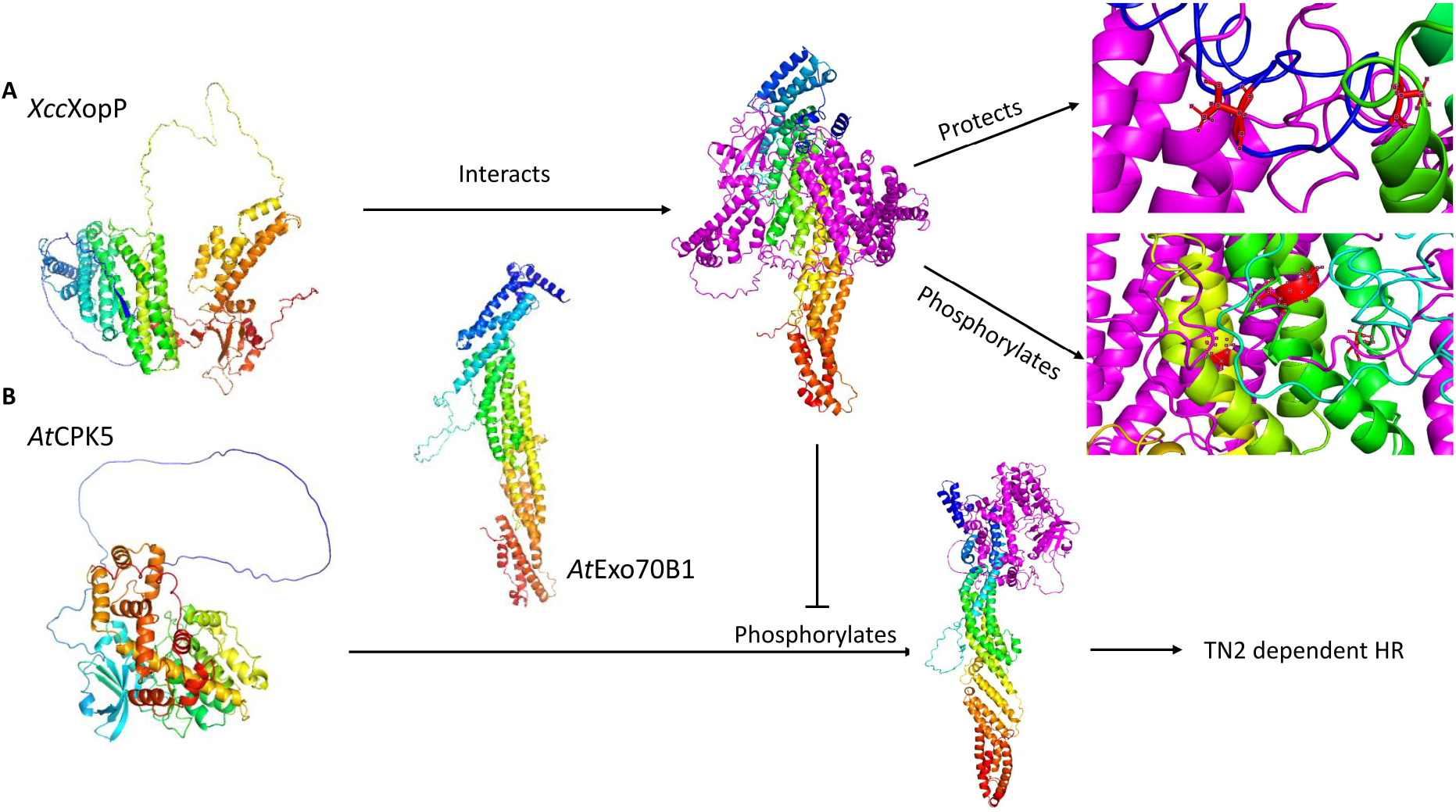
Proposed mechanism based on tertiary structure prediction. **A.** *Xcc*XopP (multicolor) interacts with *At*Exo70B1 (multicolor), forming a complex (*Xcc*XopP magenta, *At*Exo70B1 multicolor). Examples of the protected (top) and phosphorylated (bottom) residues are highlighted in red. *Xcc*XopP both phosprorylates and protects *At*Exo70B1 from AtCPK5 phosphorylation, **B**. AtCPK5 phosphorylates CPK5 and leads to TN2 dependent HR.

## Acknowledgements

We want to thank Dr. Glykeria Mermigka, Dr. Christos Christakis and Mr. Nikos Arapitsas for their advice and fruitful discussion throughout the experiments.

## Author contributions

P.F.S. and K.K. designed the research and wrote the manuscript; P.F.S. supervised the project and edited the manuscript; K.K. carried out the experiments; V.A.M. carried out *in planta* and confocal microscopy experiments; D.T. contributed lab materials; D.K., A.K., P.H.N.C. contributed material and supervised protocols; N.K. performed nanoLC-MS/MS experiments; M.K. edited the manuscript.

## Financial Disclosure Statement

K.K. and P.F.S. were supported by the European Union and Greek national funds through the Operational Program Competitiveness, Entrepreneurship and Innovation, under the call RESEARCH–CREATE–INNOVATE (“INNOVA-PROTECT” with project code: T1EDK-01878). K.K. was also supported by EMBO Short-Term Fellowship program, number 8587 and the iNEXT-Discovery, project number 871037, funded by the Horizon 2020 program of the European Commission. V.A.M. was supported by the Hellenic Foundation for Research and Innovation (HFRI) and the General Secretariat for Research and Technology (GSRT), under the HFRI PhD Fellowship grant (GA. no. 4776) and by the internal PhD supporting scheme of IMBB-FORTH.

## Supplementary Tables and Figure Legends

**Table S1.** XopP-Exo70B1 in vitro kinase assay Phosphorylated peptides

**Table S2.** CPK5-Exo70B1-XopP in vitro kinase assay

**Figure S1.** Alphafold protein structure predictions. **A.** Tertiary structure prediction of *Hordeum vulgare* subsp. *vulgare* RGH2 NLR protein (N-terminus blue, C-terminus red), **B**. Superimposition between *Hv*RGH2 (multicolor) and *At*Exo70B1 (cyan).

**Figure S2.** Structure prediction and superimposition of *Xcc*XopP homologs. **A.** *Xanthomonas oryzae* pv. *oryzae* XopP-1 (XOO_3425) superimposition (magenta), **B.** *Xanthomonas oryzae* pv. *oryzae* XopP-2 (XOO_3426) superimposition (magenta), **C.** *Xanthomonas oryzae* pv. *oryzicola* XopP-1 (XOC_1262) superimposition (magenta), **D.** *Xanthomonas oryzae* pv. *oryzicola* XopP-2 (XOC_1263) superimposition (magenta), **E.** *Ralstonia solanacearum* HLK3 (RSp0160) superimposition (magenta), **F.** *Acidovorax citrulli* XopP (Acav_0588) superimposition (magenta). *Xcc*XopP cyan. Uniprot gene entry numbers in parenthesis. Conserved β-sheet domain in frame.

**Figure S3.** Mass spectrometry imaging of phosphorylated *At*Exo70B1 peptides by *Xcc*XopP. Peptides containing residues **A.**Ser107, **B.** Ser111 residues, **C.** Ser248 residue, **D** Ser308, **E.** Thr309 and **F.** Thr364.

**Figure S4.** Mass spectrometry imaging of not phosphorylated *At*Exo70B1 peptides by *Xcc*XopP_1-420_. Peptides containing residues **A.** Ser107 and Ser111 residues, **C.** Ser248 residue, **D** Ser308 andThr309 and **D.** Thr364.

## Notes

### Competing Interest Statement

The authors have declared no competing interest.

