## Supplementary Figures for "The functional and structural characterization of *Xanthomonas campestris* pv. *campestris* core effector XopP revealed a new kinase activity"

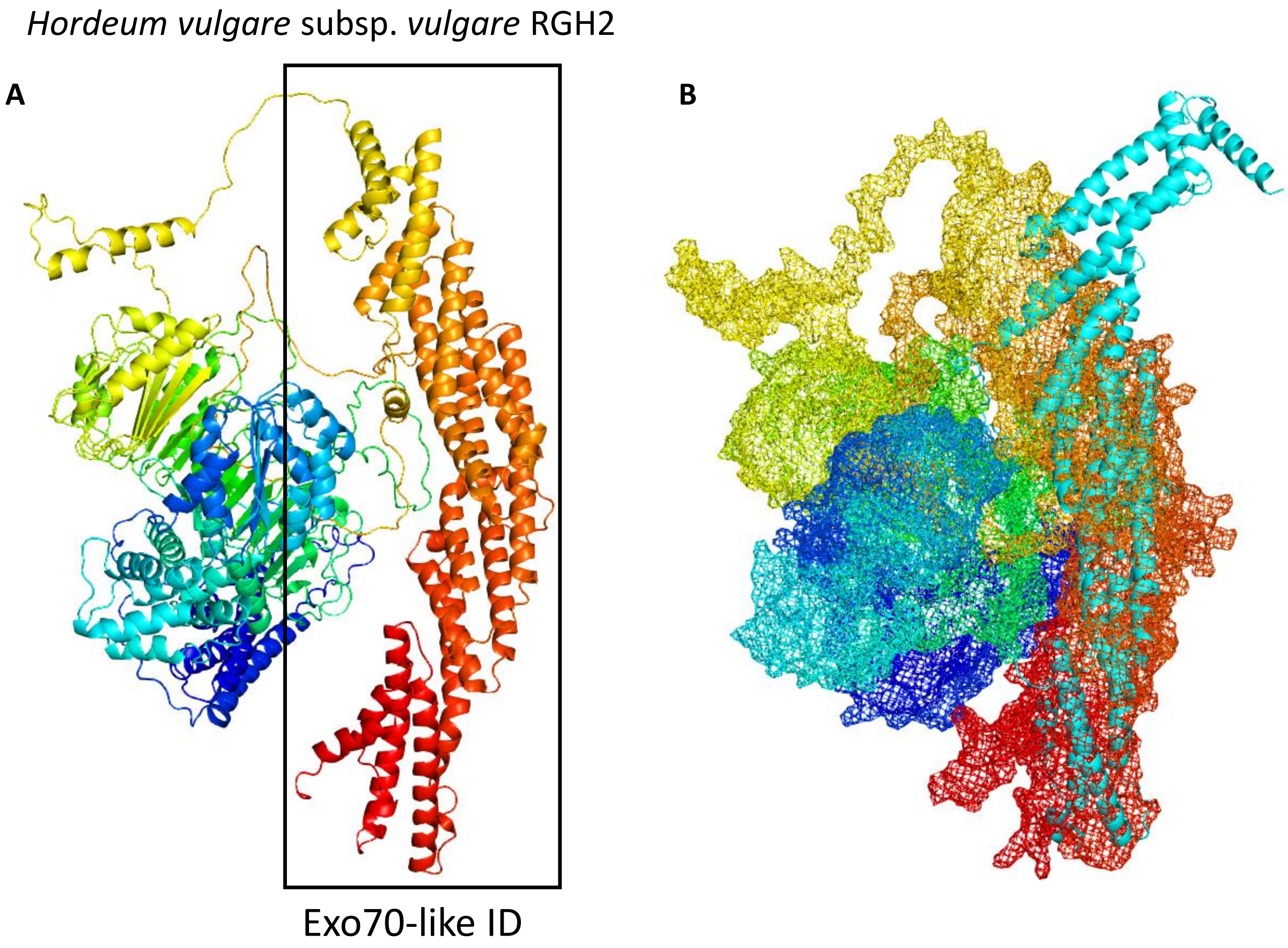


**Figure S1.** Alphafold protein structure predictions. **A.** Tertiary structure prediction of *Hordeum vulgare* subsp. *vulgare* RGH2 NLR protein (N-terminus blue, C-terminus red), **B.** Superimposition between *Hv*RGH2 (multicolor) and *At*Exo70B1 (cyan).

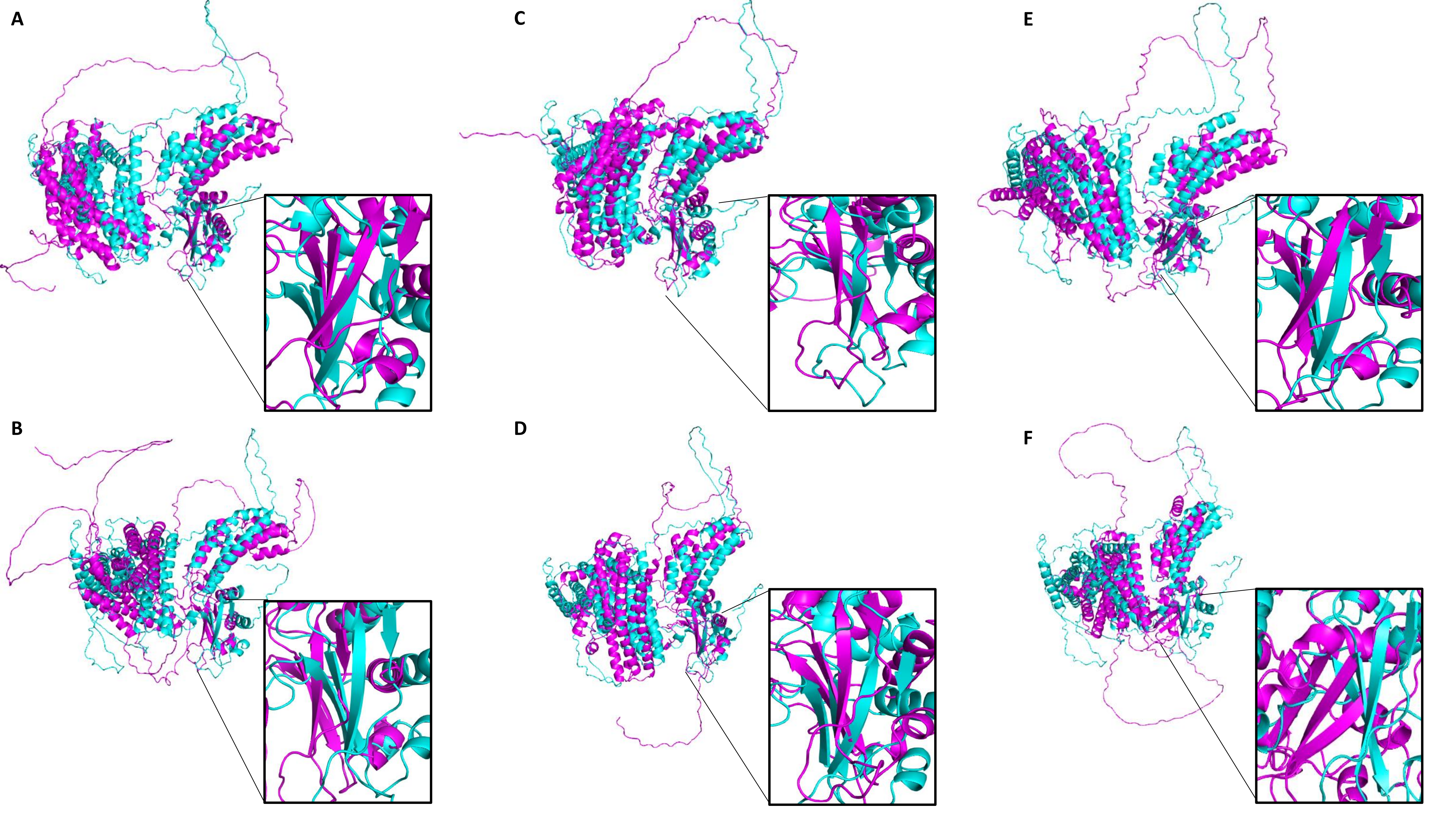
A
B
C
D
E
F

**Figure S2.** Structure prediction and superimposition of *Xcc*XopP homologs. **A.** *Xanthomonas oryzae* pv. *oryzae* XopP-1 (XOO_3425) superimposition (magenta), **B.** *Xanthomonas oryzae* pv. *oryzae* XopP-2 (XOO_3426) superimposition (magenta), **C.** *Xanthomonas oryzae* pv. *oryzicola* XopP-1 (XOC_1262) superimposition (magenta), **D.** *Xanthomonas oryzae* pv. *oryzicola* XopP-2 (XOC_1263) superimposition (magenta), **E.** *Ralstonia solanacearum* HLK3 (RSp0160) superimposition (magenta), **F.** *Acidovorax citrulli* XopP (Acav_0588) superimposition (magenta). *Xcc*XopP cyan. Uniprot gene entry numbers in parenthesis. Conserved β-sheet domain in frame.

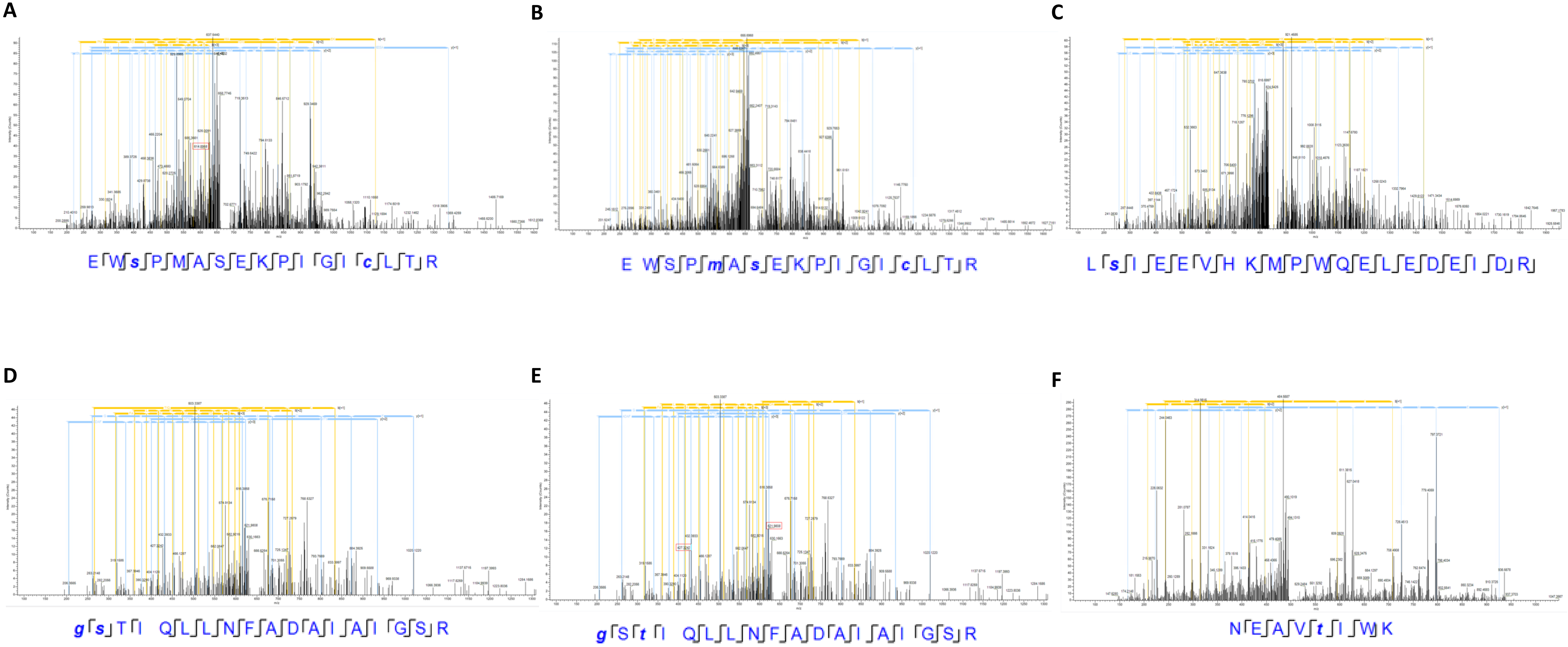


**Figure S3.** Mass spectrometry imaging of phosphorylated *At*Exo70B1 peptides by *Xcc*XopP. Peptides containing residues **A.** Ser107, **B.** Ser111 residues, **C.** Ser248 residue, **D** Ser308, **E.** Thr309 and **F.** Thr364.

**Figure S4.** Mass spectrometry imaging of not phosphorylated *At*Exo70B1 peptides by *Xcc*XopP$_{1-420}$. Peptides containing residues **A.** Ser107 and Ser111 residues, **C.** Ser248 residue, **D** Ser308 andThr309 and **D.** Thr364.

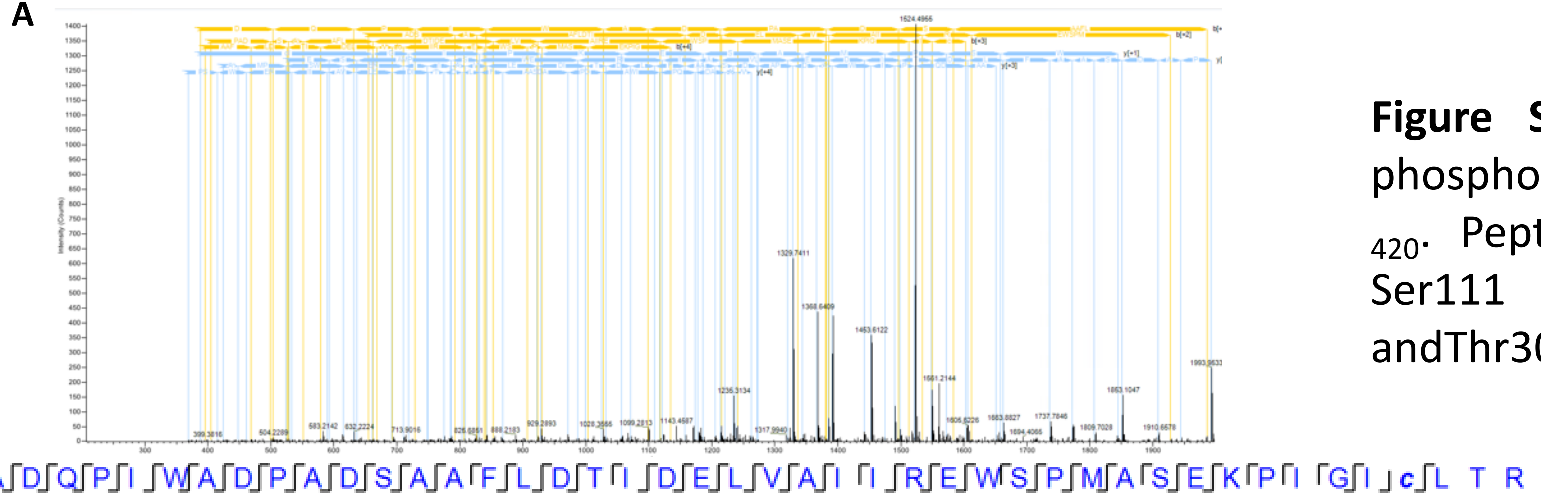

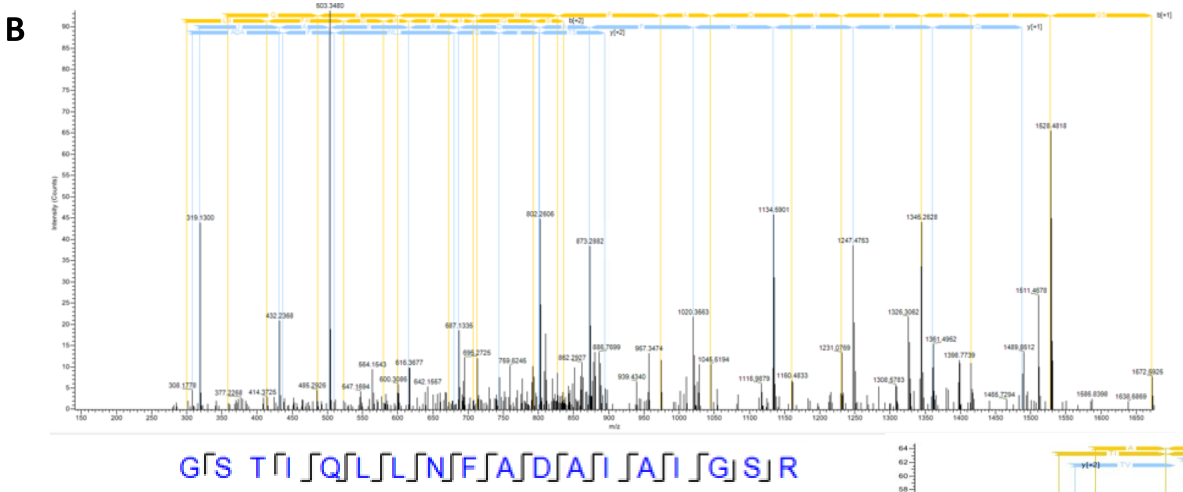

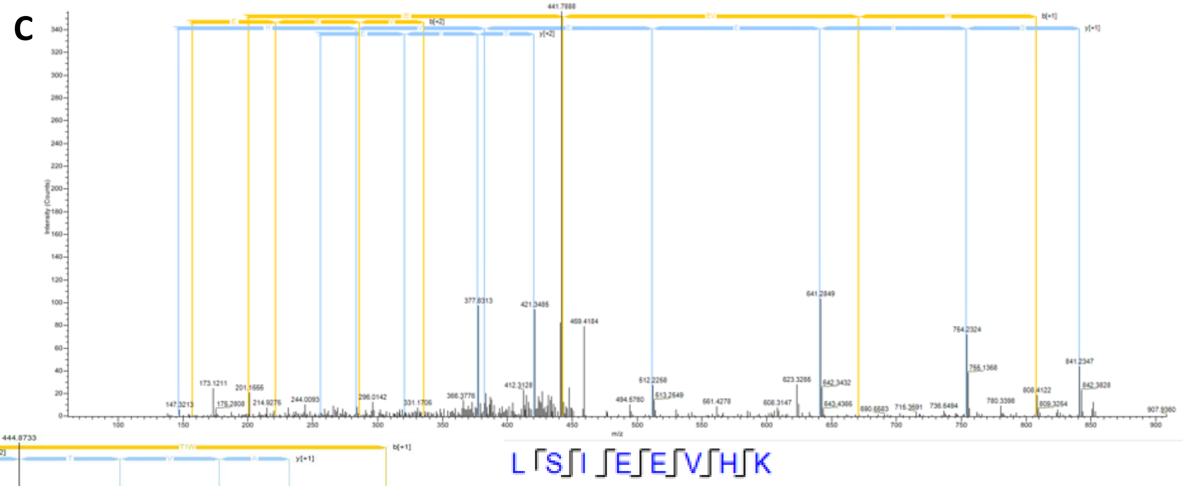

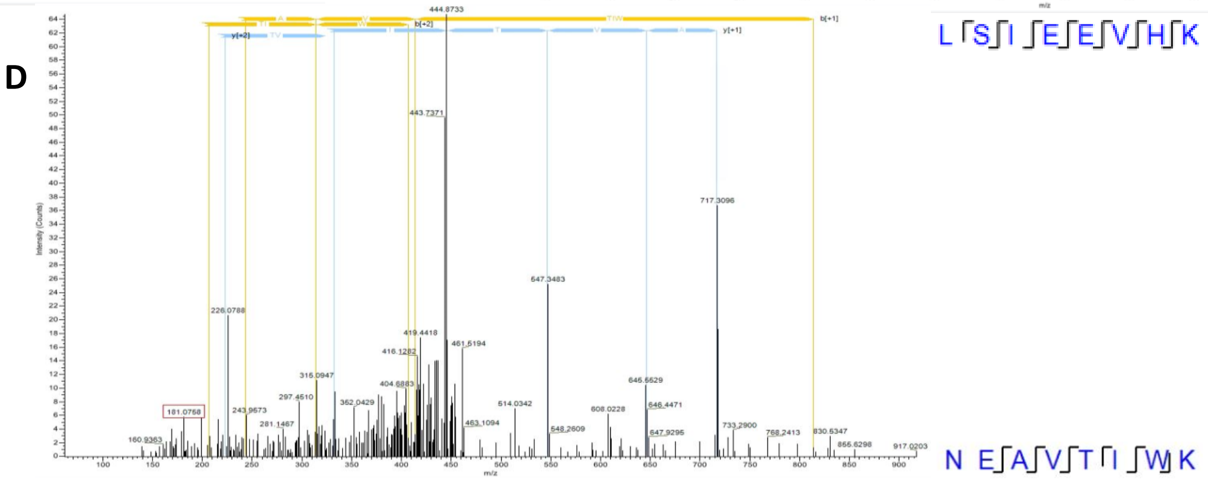
